# The functional organisation of retrosplenial feedback to V1

**DOI:** 10.1101/2025.09.25.678583

**Authors:** Melina Timplalexi, Pedro Mateos-Aparicio, William M. Connelly, Adam Ranson

**Author notes:** Corresponding author: Adam Ranson.

## Abstract

Cortical feedback from higher frontal and association areas to early sensory cortex is thought to contribute to a range of cognitive processes in which sensory signals are processed in a context sensitive manner. Despite the ubiquity of feedback circuitry in the cortex, the information carried by feedback connections, and the principles governing how it is targeted from area to area remains poorly understood. Here we characterized the functional properties of a prominent feedback circuit which links retrosplenial cortex (RSC) and primary visual cortex (V1). We found that RSC→V1 axons relay retinotopically selective signals to V1 which at a coarse scale match the region of the V1 retinotopic map innervated, were tuned for spatial and temporal frequency, but not tuned for orientation. Two-color imaging of RSC→V1 boutons and local L2/3 V1 neurons further revealed that at a finer scale RSC→V1 bouton receptive fields are systematically offset relative to those of V1 in the nasal direction, consistent with RSC→V1 boutons conveying signals predictive of upcoming V1 activity in the retinotopic region innervated during forward movement.

## Introduction

The cortex is constituted of multiple hierarchically organised areas which interact through feedforward and feedback pathways to produce progressively more abstract representations. This organisation permits adaptive processing of sensory information and extraction of visual features to support sensory guided cognitive processes. Feedback pathways from ‘higher’ cortical areas encoding more abstract sensory and non-sensory information to early sensory areas are thought to be critical to this process of adaptive sensory encoding. Feedback circuitry has been proposed to serve diverse roles, including relaying predictions, motor and attentional signals to bias or transform representations in lower sensory areas. Our understanding of what information is relayed in feedback signals, and how they are targeted is limited.

One relatively well understood short range feedback circuit in the mouse cortex is that linking the higher visual area LM to V1 with which it is reciprocally connected. Here a dense feedback projection relays grossly retinotopically matched sensory signals representing an area in visual space translated along the axis of the LM axon’s preferred direction to neurons in V1 (*1, 2*) and to act via recruitment of nonlinear processes in tuft dendrites of V1 pyramidal neurons (*3*). Other experiments have provided evidence that such feedback is responsible for generation of a second receptive field of V1 neurons which is mutually antagonistic with the primary feedforward receptive field (*4*). Broadly speaking, the proposed computational role of such an arrangement is that it would allow the relaying of predictions to V1 which may enhance or supress expected sensory driven neural responses (*1–4*).

Another longer range circuit in mice monosynaptically connects medial frontal cortical regions (including areas referred to as anterior cingulate cortex, A24b and M2) and the primary visual cortex (*5*). This projection has been shown to be in part visual and motor driven (*6*) and has been demonstrated to exert a retinotopically selective influence on sensory processing (*5*) which share similarities with some forms of selective visual attention described in primates (*7– 10*), although the manner in which this circuit is endogenously recruited remains unclear (*5, 6, 11–13*).

Here we examine the largely uncharacterised functional organisation of feedback to V1 from the retrosplenial cortex (RSC) - an intermediate associational area which is reciprocally connected to the hippocampal formation, anterior thalamic nuclei and provides one of the densest sources of feedback to V1 (*14*). RSC has been implicated in diverse processes associated with spatial memory and navigation (*15*). The dysgranular RSC possesses a coarse-grained retinotopic map (*16*) and contains a fraction of neurons selective to orientation (*16, 17*). We used two-photon imaging of RSC→V1 axon terminals to show that RSC relays low spatial and temporal resolution visual feedback signals to V1 which are retinotopically selective and targeted to approximately spatially matched areas of the V1 retinotopic map encoding the same area of visual field. At finer scale RSC→V1 input retinotopy was found to be slightly spatially offset in the nasal direction relative to the region of V1 retinotopic map targeted. This offset was most pronounced in the monocular visual space directly lateral to the mouse’s head, where visual flow during forward locomotion is fastest, and where visual predictions provided by feedback may therefore need to be most spatially offset on average.

## Results

### RSC relays spatially selective feedback to primary visual cortex

We first aimed to confirm the presence of the previously reported anatomical feedback connection from dysgranular layers of the retrosplenial cortex to the primary visual cortex (*14, 18*). To visualise RSC→V1 axons we labelled them with axon targeted jGCaMP8s by injecting an AAV into layer 5 of the retrosplenial cortex (**Figure 1A**) and then visualised axons of expressing RSC neurons in primary visual cortex (**Figure 1B**). Consistent with previous reports, we observed RSC afferents distributed throughout the depth of V1, with a notable concentration in layer 1 (**Figure 1B**).

**Figure 1.**
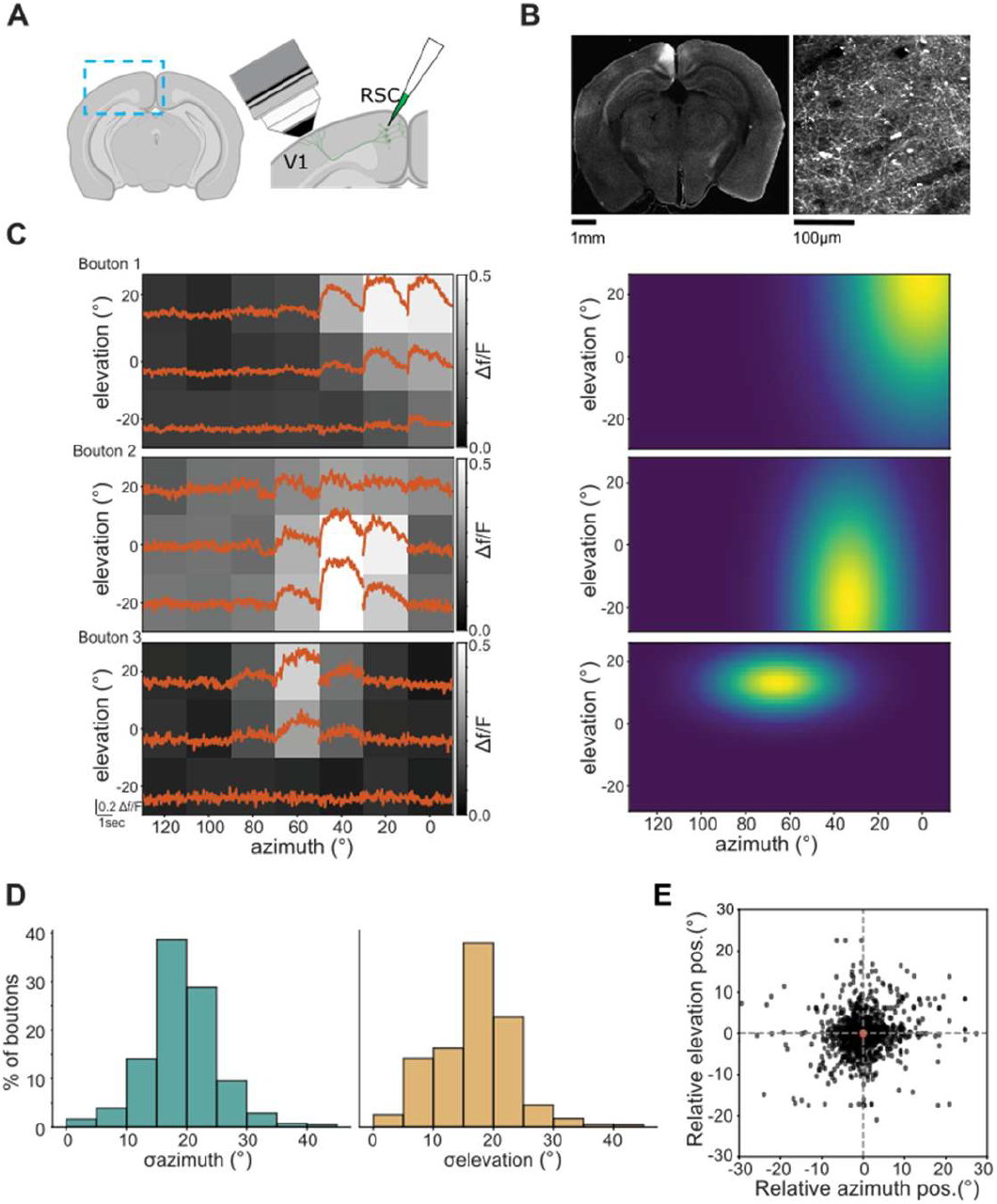
RSC→V1 boutons are retinotopically tuned and co-tuned boutons are clustered. (A) Schematic of injection for labelling of RSC→V1 axons and two-photon recording. (B) Example field of view showing RSC axons in layer 1 of V1 labelled with jGCaMP8s. (C) Example spatial responses (average traces plotted over heatmap) for three visually responsive boutons. (D) 2d Gaussian fits of the spatial receptive fields of boutons shown in C. (E) Distribution of spatial receptive field size (sigma from fit) in azimuth and elevation. (F) Distribution of fitted preferred position of each RSC→V1 bouton relative to the preferred position of other boutons in the same field of view. n = 1950 boutons collected from 8 mice in 12 experiments.

**Figure 2.**
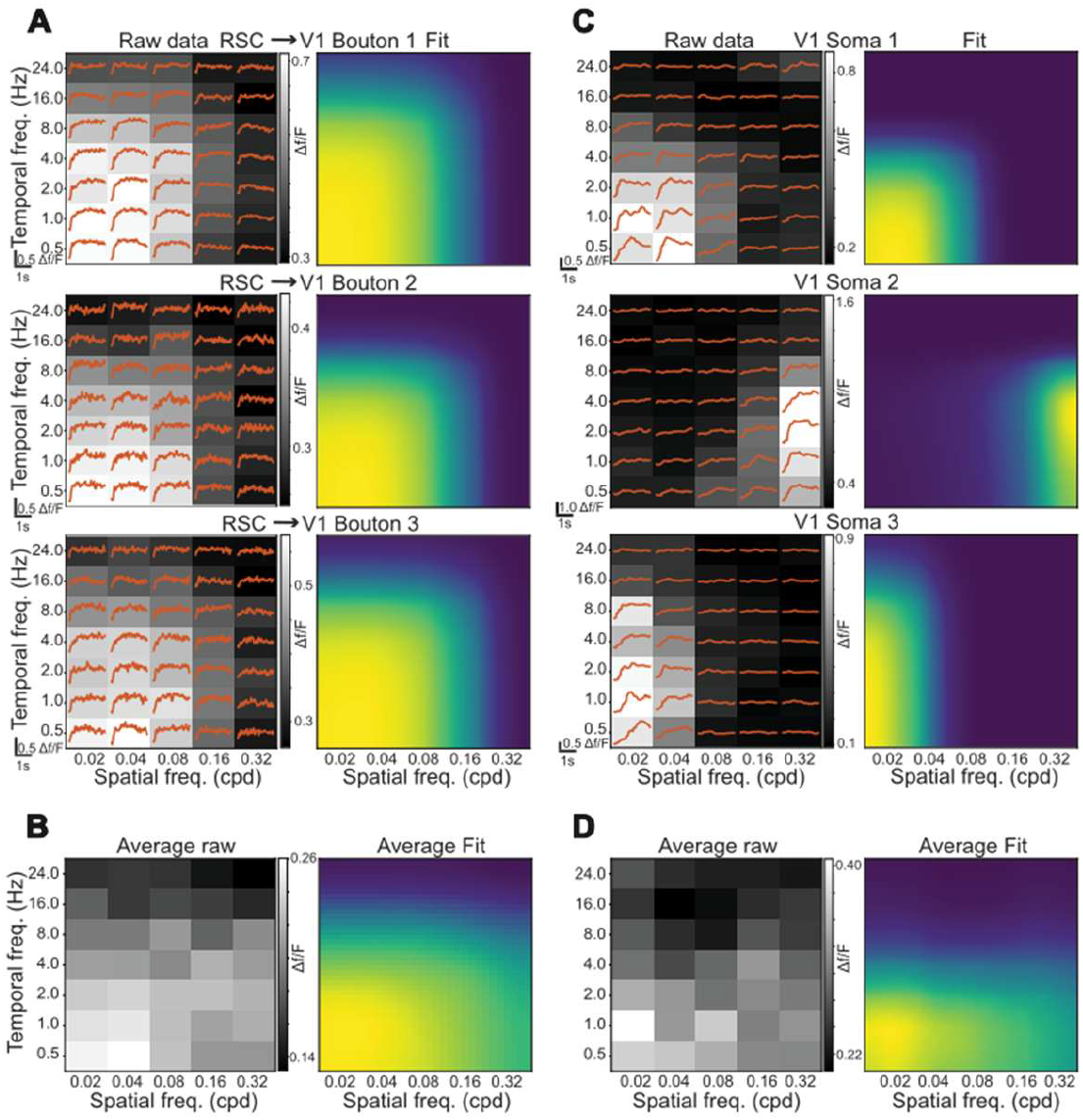
RSC→V1 axons relay temporal and spatial frequency tuned signals. (A) Example responses to different spatial and temporal frequencies (average traces plotted over heatmap) for three visually responsive RSC→V1 boutons (left), and 2d Gaussian fits of spatial and temporal frequency tuning of the same boutons. (B) Average responses of population of RSC→V1 boutons. (C-D) As in A-B, but for V1 somas. For V1 soma recordings, n = 13101 neurons from 4 mice from 4 experiments. For RSC→V1 bouton recordings, n = 1035 boutons from 7 recordings in 7 mice.

We next examined the spatial response selectivity of RSC→V1 boutons. Retinotopically selectivity has previously been reported in the responses of many RSC somas (*16*) and so we first asked whether RSC→V1 feedback signals are also retinotopically selective. To test for retinotopic selectivity, awake head-fixed mice were presented with circular 30° horizontally oriented drifting grating stimuli located in a 7 x 3 grid with 20° spacing. We found that 69 ±27% of boutons were visually responsive to stimulation in at least one position and that responsive boutons tended to show strong spatial selectivity (**Figure 1C**). We quantified average response amplitudes of RSC→V1 boutons to each retinotopic position and fit a 2d Gaussian to the responses of each bouton such that the σ_azimuth_ and σ_elevation_ indicate RSC→V1 bouton receptive field extent (**Figure 1C right**). Since boutons tended to respond to more than one retinotopic location, the fit allowed more fine scale estimation of receptive field centre despite the relatively course mapping stimulus spacing. On average we found that RSC→V1 boutons were spatially tuned (**Figure 1D**, azimuth mean σ = 19.87 ± 5.47; elevation mean σ = 16.99 ± 6.57) and that within each field of view, fitted receptive field centres tended to be similar (mean absolute difference of fitted receptive field centre from average of field of view = 7.24 ± 2.21 **Figure 1E**). These findings indicate that RSC→V1 axons feedback visual information about confined regions of the visual field to primary visual cortex, and the clustering of spatial selectivity within a field of view is suggestive of a retinotopic organisation of feedback axons relative the V1 retinotopic map as has been reported in feedback to V1 from higher visual area LM (*1, 2*). Retrosplenial feedback to V1 is thus well placed to provide spatially targeted top-down modulation of V1 activity.

### RSC→V1 axons are spatial and temporal frequency tuned but not selective to orientation

In order to better understand the form of modulation RSC→V1 axonal feedback may be exerting on V1 sensory processing, we examined the spatial and temporal frequency tuning of RSC→V1 axons. We hypothesised that relative to the broad range of spatial and temporal frequency selectivity in V1, RSC→V1 feedback may exert modulatory influence at lower temporal frequencies in accordance with the role of RSC in processing distal landmarks. Consistent with this, RSC boutons were preferentially tuned to lower temporal frequencies than V1 neurons recorded in a separate cohort of mice while mean spatial frequency tuning was similar (**Figure 1A-D;** RSC TF peak = 1.46 Hz, V1 peak = 2.88 Hz; t-test p = 1e^-20^; RSC SF peak = 0.12 cpd, V1 SF peak = 0.11 cpd; t-test p = 0.3). We next examined more closely the selectivity of visual feedback signals to V1 to determine if they more likely relay a broad spatial modulatory signal or more detailed predictive information to V1. A subset of somas of neurons in RSC have previously been shown to be orientation selective (*16, 17*) and so we tested if RSC feeds back orientation specific signals to V1. Responses of RSC boutons to gratings (aligned to the preferred retinotopic location of the field of view of boutons) showed almost no orientation selectivity (**Figure 3A-C**) indicating that it is the subset of non-orientation tuned RSC neurons which provide feedback to V1, and that RSC is unlikely to be feeding back fine-grained visual predictive signals to V1.

**Figure 3.**
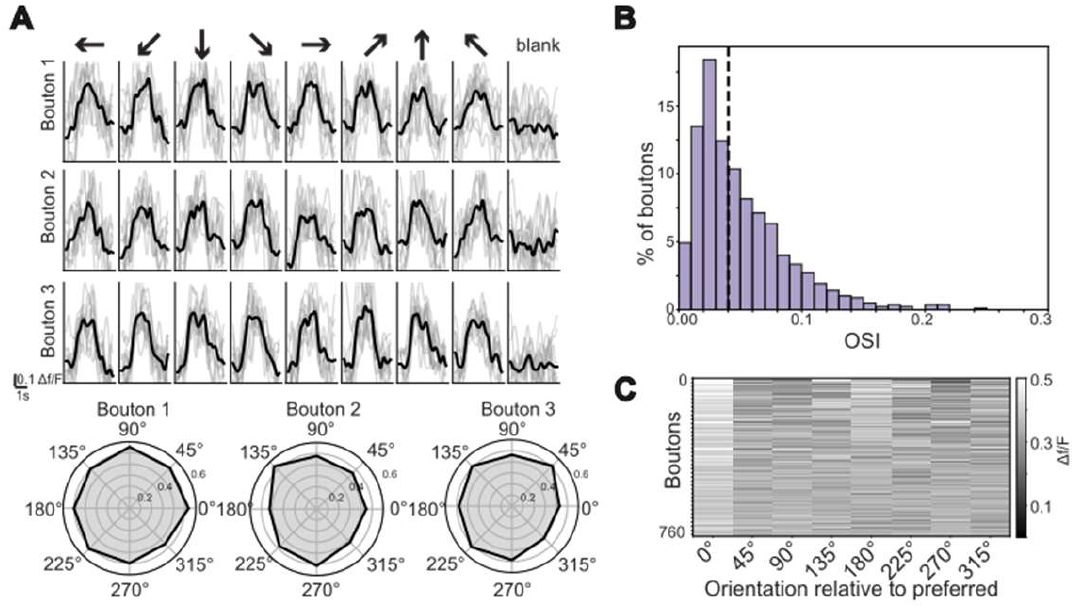
RSC→V1 boutons are not tuned to orientation. (A) responses of 3 **stimulus** Example RSC→V1 boutons to different grating orientations/directions and blank (upper), and polar plots of tuning (lower). (B) Distribution of OSI across whole population of characterised boutons. (C) Orientation responses of RSC→V1 boutons relative to each bouton’s preferred orientation. n = 827 boutons from 11 recording in 7 mice.

### RSC→V1 axons target retinotopically matched regions of V1

Given the retinotopic selectivity of RSC→V1 boutons, and the similarity of retinotopic tuning of boutons observed within each field of view we next asked how bouton retinotopic tuning relates to the retinotopic tuning of the V1 retinotopic map area innervated. To test this, mice were co-labelled with the genetically encoded calcium indicators jRGECO1a (in V1 neurons; **Figure 4A**, red) and jGCaMP8s-axon (in RSC→V1 axons; **Figure 4A**, green). This approach allowed measurement of retinotopy of RSC→V1 axons in layer 1 and layer 2/3 V1 somas in the same mice (**Figure 4B**). Mice were stimulated with the retinotopic mapping stimulus described above, and spatial selectivity was again determined by fitting the mean responses of each RSC→V1 bouton or V1 soma using a 2D Gaussian. For each field of view, we then calculated the median azimuth and elevation preference of each RSC bouton and of the local V1 somas. Although highly scattered, RSC bouton receptive field centres were on average matched to somas in the region of V1 they targeted (mean absolute difference = 4.37 ± 4.41) although a fraction of boutons conveyed visual signals from more distal visual field locations (**Figure 4D**).

**Figure 4.**
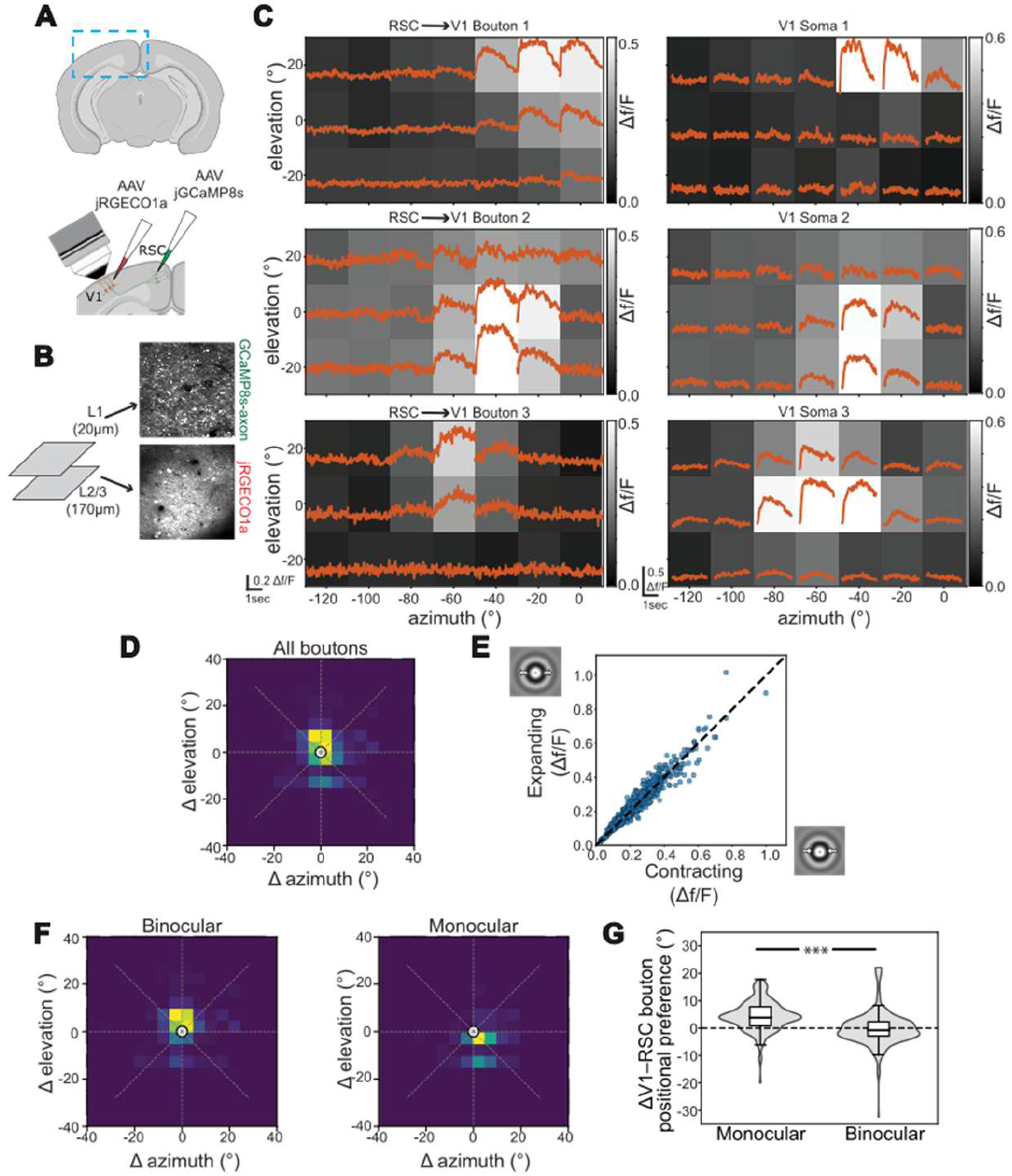
Alignment of RSC→V1 bouton spatial tuning relative to targeted region of V1. (A) Schematic of injection for labelling of V1 somas with jRGECO1a and RSC→V1 axons with jGCaMP8s and two-photon recording. (B) Example field of view showing RSC axons in layer 1 of V1 labelled with jGCaMP8s and layer 2/3 somas labelled with jRGECO1a. (C) Example spatial responses (average traces plotted over heatmap) for three visually responsive boutons (left) and 3 retinotopically aligned V1 somas (right). (D) Heat plot showing distribution of difference in tuning of all RSC→V1 boutons relative to local V1 neurons in azimuth and elevation. (E) Responses of each RSC→V1 bouton to expanding versus contracting stimulus. (F) Data from D divided into binocular and monocular areas of the V1 retinotopic map showing nasal offset of RSC→V1 bouton positional tuning relative to V1 tuning in monocular area. (G) Distribution of ΔV1-RSC bouton positional preference showing offset is specific to monocular visual space. For V1 soma recordings, n = 827 neurons from 9 experiments in 6 mice. For RSC→V1 bouton recordings, n = 1467 boutons from 9 experiments in the same 6 mice used for soma recordings.

Previous studies of shorter range visual feedback from LM to V1 have reported a systematic offset in retinotopic targeting such that direction-selective axons targeted V1 retinotopic sites which are orthogonal to the tuning of the feedback axon (*1, 2*). Since we observed no direction selectivity in RSC boutons, such an arrangement would not be possible in this case, but we reasoned that boutons might instead possess tuning more generally for stimuli which would be expected to proceed on average activation of the targeted V1 site. To test for this, we examined responses to expanding or contracting radial grating stimuli, centred on the average field of view retinotopically preferred location, and hypothesised boutons would exhibit a selectivity to contracting stimuli. However, while RSC boutons did respond to both contracting and expanding radial grating stimuli, they exhibited no preference for radial flow direction (**Figure 4E**). Closer inspection of differences in azimuth tuning between RSC boutons and V1 neurons (Δazimuth) in the same field of view showed a small systematic offset in position tuning such that RSC boutons on average encoded a more nasal retinotopic position relative to V1 neurons within the retinotopic region innervated (**Figure F**). We therefore next hypothesized that if RSC→V1 axons are relaying predictive signals to V1 which anticipate activity in the part of the retinotopic map they target, then differences in velocity of visual flow lateral to the animal as compared to in front of the animal (i.e. during forward locomotion) may result in variation in the magnitude of Δazimuth depending upon visual field location. Specifically, Δazimuth might be expected to be larger in monocular areas of the visual field which are subject to higher velocity visual flow during forward movement. Consistent with this possibility, we found that the observed non-zero Δazimuth was driven by boutons encoding monocular visual space and targeting monocular V1. While mean Δazimuth for boutons encoding the more nasal 0° to 30° of visual space (i.e. binocular) was close to zero (0.01° ± 0.15), mean Δazimuth for boutons encoding the more temporal 30° to 90° of visual space was 5.82° ± 0.50 (Mann–Whitney U two-sided test, p = 1e^-90^; **Figure 4F and G**). In summary, on average monocular visual field encoding RSC→V1 boutons exhibit a systematic nasal-azimuth offset in their spatial tuning relative to the V1 neurons in the region of the V1 retinotopic map they target.

### Discussion

Here we use two-photon imaging of RSC boutons during visual stimulation to provide the first detailed characterisation of visual feedback signals relayed from the retrosplenial cortex to the primary visual cortex in awake viewing mice. We found that RSC→V1 axons feedback spatially specific signals to V1 from regions of approximately 20-30° of visual space. The RSC→V1 bouton population as a whole was found to differ notably in its temporal frequency tuning relative to neurons in V1 from which RSC receives direct input (*19*). Specifically, RSC axons were found to feedback relatively low temporal frequency signals to V1. This suggests a potential role in modulation of V1 visual representations which share this features such as distal visual landmarks which are known to be encoded by neurons within RSC (*19*). In contrast to reports from RSC somas, we found that the subpopulation of RSC neurons from which RSC→V1 axons originate were not selective to grating stimulus orientation. In this sense visual RSC feedback differs from that from the higher visual area LM which feeds back orientation tuned signals to V1 (*2*). Dual-color imaging of RSC→V1 axons and V1 somas revealed that while at a coarse scale visually driven feedback is on average retinotopically targeted to spatially matched areas of the V1 retinotopic map, at finer scale RSC→V1 feedback is retinotopically slightly offset in the nasal direction relative to local V1 neurons. Interestingly, this effect is specific to the monocular visual field, which may be due to differences in velocity of monocular versus binocular visual flow experienced during forward locomotion. This organisation shares similarities with feedback from LM to V1 which is spatially offset across the visual field in a direction which dependents on bouton directional tuning. This similarity suggests a simple general connectivity rule based on temporal correlation may be more generally predictive of feedback connectivity.

Retrosplenial cortex is therefore well placed to spatially modulate visual processing in a top-down manner and may provide visual predictions to V1 of upcoming visual input during forward movement. This could prime V1 neurons in a given region to be more responsive when the animal expects a landmark or other visual cue to appear in that location. The recruitment of the circuit described here could alternatively represent a form of stimulus driven bottom-up or exogenous spatial attention. It would therefore be of interest to determine if this circuit can be recruited volitionally to selectively modulate or enhance processing of specific regions of the visual field depending on task demands as has been shown in selective spatial attention paradigms in mice (*20*).

When relaying feedback in the ‘stimulus driven’ regime explored here, the lack of orientation tuning of RSC→V1 boutons suggests that RSC feedback provides spatial gain modulation rather than precise feature-specific input. It will be of interest to determine whether this lack of tuning is also observed in a context of more active vision such as during visual discrimination or navigation. The one previous study of activity of RSC→V1 bouton activity measured activation during visually guided associative learning task and provided evidence for a shift in the balance from bottom-up to top-down (i.e. from RSC) ‘information streams’ (*18*). However, under the stimulus driven regime explored here, the RSC feedback stream does not encode the visual feature being discriminated in the task employed in this study (i.e. grating orientation). This suggests that when recruited through internal drivers (rather than driven by external stimuli) this circuit may be capable of a greater degree of selectivity, perhaps through activation of sparse subsets of RSC→V1 projecting neurons, or through a higher degree of selectivity in postsynaptic targeting in V1. An alternative explanation of the apparent lack of visual feature tuning we observe is that RSC boutons have complex receptive fields which are not well discriminated using grating stimuli, and it would therefore be of interest to probe RSC bouton receptive fields with a natural or sparse noise stimulus.

We observed a significant fraction of RSC→V1 axons which we were unable to drive with the visual stimuli used here, prompting the question of the role they serve. One likely possibility is that we did not provide suitable visual stimuli to activate these neurons. Stimulation with a more expansive selection of natural video stimuli will help determine this. Another possibility is that RSC→V1 boutons encode some behavioural or other internal state, or conjugation of states not examined in this study.

The computational function conferred by RSC→V1 feedback depends upon postsynaptic partners of RSC→V1 boutons. Feedback to primary visual cortex has been shown to target somatostatin-expressing interneurons (*5, 21*), VIP-expressing interneurons (*5*), and excitatory neurons (*5, 18*). The non-specificity of visual feature tuning observed in RSC boutons may be an indication that post synaptic partners are similarly broadly tuned, suggesting potential interneuron targeting (*22*). Determining at a functional and cell-type level how RSC→V1 boutons are paired to postsynaptic V1 neurons will be critical to determine whether this feedback circuit has the effect of, for example, supressing or facilitating expected visual input and inferring what computational function this feedback circuit serves.

In summary this study provides a first detailed description of the visual tuning properties of RSC→V1 feedback axons and their relationship to tuning properties of local V1 neurons in regions targeted, and therefore a basis on which future work can examine how this circuit is recruited during active sensation to allow context sensitive processing of visual information.

## Limitations of the study

The study provides the first detailed characterisation of RSC→V1 feedback circuit organisation and visual responses, however deeper study of visual responses to natural stimuli would help further confirm the lack of visual tuning of RSC→V1 boutons to detailed spatial features. Relatedly, we limited experiments to passive viewing of sensory stimuli, and recruitment of these circuits may differ in important ways during visually driven behaviour such as during navigation, or attentive visual discrimination. A second limitation is that we confined our analysis to RSC→V1 boutons in the most superficial layers of V1, while we did observe boutons throughout the depth of V1. This may bias our sample of boutons to those synapsing onto superficial post synaptic structures such as L1 interneurons and tuft dendrites. Tuning properties of deeper synapsing RSC→V1 boutons may therefore differ from those described.

## Acknowledgements

This work was supported by grants PDI 2019-109285GA-I00, RYC2021-032313-I, PID2022-139822NB-I00, PCI2022-134995-2 to A.R and PRE2020-093128 to M.T. from the Spanish Secretary of Research, Development and Innovation (MINECO), and has received funding from the European Union’s Horizon ERC Grants research and innovation programme under ERC Consolidator grant agreement No. 101088598 (HEU-101088598-DREAMNET) to A.R.

## Experimental model and subject details

### Animals

All experimental procedures were carried out under the Ethics Committee on Animal and Human Experimentation from the Universitat Autònoma de Barcelona and followed the European Communities Council Directive 2010/63/EU. Experiments were performed on adult (age>P90) C57BL/6J mice. Experiments were carried out in mice of either sex, group-housed, with 12:12h light-dark conditions, *ad libitum* access to food and water. All recordings were made during the light period.

## Method details

### Animal Surgical Preparation and virus injection

All surgical procedures were conducted under aseptic conditions. Before the cranial window surgery, animals were administered the anti-inflammatory drugs Rimadyl (5 mg/kg, s.c.) and dexamethasone (0.15 mg/kg, i.m.), and antibiotic Baytril (5 mg/kg, s.c.). Anesthesia was delivered in gas form with a 5% concentration of isoflurane in oxygen and then maintained at 1.5-2% for the duration of the surgery. Once anaesthetised, mice were placed onto a heating pad to maintain their body temperature at 37°C, and head fixed using ear bars into a stereotaxic frame. The area of the surgery was shaved and cleaned using a 70% ethanol solution followed by an antiseptic iodine solution, and two injections of lidocaine (2% w/v) were made subcutaneously into the scalp for local anaesthesia. The scalp and the periosteum overlying the area of headplate fixation were then resected.

For RSC injection, a small craniotomy was made above the retrosplenial cortex (-2.8mm posterior from bregma, 0.3mm lateral from the midline) using a dental drill, thinning of the bone and then removal of a small bone flap using a hypodermic needle. After the brain was exposed, an AAV was injected at a depth of 400µm to drive expression of jGCaMP8s (pAAV-hSynapsin1-axon-jGCaMP8s-P2A-mRuby3, Addgene plasmid #172921, 5 × 10^12^ GC per millilitre titre, volume 100 nL). The craniotomy of the RSC was sealed with Vetbond. A custom head plate was then attached to the skull using Vetbond and secured with dental cement (Paladur, Heraeus Kulzer), with the aperture centered over the binocular area of the right hemisphere V1 (-3.2 posterior and 3.1 lateral from bregma). A 3 mm circular craniotomy was next performed, centered on the stereotaxically identified binocular area of V1.

For experiments in which V1 neurons were labelled, after the V1 craniotomy was performed, an AAV was injected at multiple sites at a depth of 200-400µm to drive expression of jGCaMP7f (pGP-AAV-syn-jGCaMP7f-WPRE, Addgene plasmid #104488, 5 × 10^12^ GC per millilitre titre, volume 100 nL) or jRGECO1a (AAV1.Syn.NES-jRGECO1a.WPRE.SV40, Addgene plasmid #100854, 10 × 10^12^ GC per millilitre titre, volume 100 nl, used in experiments were both axons from RSC and V1 neurons were recorded).

Injections were made using either a Nanoject II system (Drummond Scientific Company) or a Stereodrive Drill & Microinjection Robot (Neurostar) at a rate of 10 nL/min using pulled and beveled oil-filled glass micropipettes with a tip outer diameter of approximately 30-40. After injections, the craniotomy over V1 was closed with a glass insert constructed from 3 layers of circular glass 100 µm thickness (1 of 4 mm and 2 of 3 mm diameter) bonded together with UV cured optical adhesive (Norland Products; catalog no. 7106). Habituation to head fixation and imaging started a minimum of 2 weeks after surgery.

### *In vivo* imaging

*In vivo* two-photon imaging was performed using a Movable Objective Microscope with Resonant Scan box (Sutter Instrument) with a N16XLWD-PF objective (Nikon, 0.8 NA, 3.0 mm WD). Imaging of the green calcium indicators jGCaMP7 and jGCaMP8 was performed by excitation at 920nm using either an 80 MHz Ti-Sapphire laser (Chameleon Ultra II, Coherent) or a 920 nm 80 MHz fiber laser (FemtoFiber ultra 920, TOPTICA). Imaging of jRGECO1a was performed by excitation at 1050nm using an 80 MHz fiber laser (FemtoFiber ultra 1050nm, TOPTICA). In all experiments laser power at the sample was limited to approximately 60mW. Data were acquired at a frame rate of 30 Hz (512 × 512 pixels) using Scanimage 5 with a standalone NI FLEX RIO DAQ (National Instruments). The field of view size in experiments in which V1 somas were recorded was approximately 250 x 250 µm^2^ while in experiments in which boutons were recorded the field of view was approximately 100 x 100 µm^2^. Synchronisation of imaging with visual stimulation and other data (behavioral and eye cameras, rotary encoder of linear treadmill) was attained by acquiring timing signals on a separate data acquisition card (NI PCIe-6323, National Instruments) using custom MATLAB code (https://github.com/ransona/Timeline-UAB).

### Behavioural and pupil position monitoring

In all the experiments videos of both eyes of the animals were recorded using USB monochrome cameras (Imaging Source model DMK 22AUC03 with lens Azure-7524 mm and a 800nm short pass filter) under IR illumination, with frames acquired using custom Python code (https://github.com/ransona/py_eye) and synchronised using a digital signal produced by an Arduino Uno (Arduino). Eye videos were processed to obtain pupil position by first detecting a number of eye landmarks using DeepLabCut (*23*). Namely, four points were detected which corresponded to the centre of upper and lower eye lid and medial and lateral eye corners, and eight points were detected which corresponded to the edges of the pupil (i.e. an octagon). The model was trained on 600 hand labelled frames and was confirmed to function well by manual inspection of experiments with detected features overlaid on eye video frames. While the network’s placement was visually indistinguishable from human placement for the vast majority of frames, the network would always attempt to place all eight pupil markers, even if the mouse was blinking or the pupil was otherwise mostly obscured by the eyelids. To remedy this, a program was written to construct an eye shape by passing a pair of parabolic curves through the medial, superior, and lateral and medial, inferior, and lateral eye markers respectively, and any pupil markers that fell outside this eye shape were discarded. In addition, if a set of basic assumptions about the eye shape were violated – for instance, if the medial eye marker was lateral to the lateral eye marker – all pupil markers were discarded. Finally, if six or more pupil markers were valid, an ellipse was fitted to them to minimize the least-squared error. Together these measures resulted in high fidelity tracking of pupil behavior across frames (*6*). Timepoints at which the pupil could not be detected were excluded from further analysis. Animal locomotion on the single axis treadmill was measured using a rotary encoder with 1024 steps per rotation (Kübler, 05.2400.1122.0100) acquired using an NI PCIe-6323 DAQ (National Instruments).

### Visual stimuli

Visual stimuli were generated using a custom written interactive visual stimulation rig (https://github.com/neurogears/ranson-rig) controllable by OSC and built upon Bonsai and BonVision (*24*). Stimuli were presented on three 54 x 40 cm^2^ LED Monitors at 60 Hz refresh rate (D24-20, Lenovo) positioned such that stimuli could be presented from approximately - 120° to +120° in azimuth, and -30° to +30° in elevation. Stimuli were warped to correct for screen flatness using Bonvision and screen luminance was linearised. To avoid contamination of imaging data by stimulus screen light, LED screens were modified in house to flicker the screen LED in synchrony with microscope line scanning such that LEDs were only illuminated during scan line turn around. A custom designed circuit was used to allow control over screen LED on time and maintenance of screen brightness during frame flyback (https://github.com/ransona/screen-LED-blanking-circuit). Stimulus timing was synchronized to other data acquisition using both digital signals produced by Bonsai via a Harp Behavior board (OEPS-1216, open ephys) and a photodiode (OEPS-3011, open ephys) which acquired an on-screen black/white alternating sync square.

Retinotopic mapping stimuli consisted of 30 x 30° circular horizontally oriented square wave gratings of spatial frequency 0.05 cycles per degree which drifted at a temporal frequency of 2 Hz. Stimuli were presented in positions on a 7 x 3 grid with stimuli centred in positions ranging from 120° to 0° in azimuth and -20° to 20° in elevation. Each grating stimulus was presented for 2 seconds with a 1 second interstimulus interval and each position was repeated 10 times in a pseudorandom order.

Stimuli to characterise orientation tuning consisted of 30 x 30° circular horizontally oriented square wave gratings of spatial frequency 0.05 cycles per degree which drifted at a temporal frequency of 2 Hz, as well as a blank stimulus. Grating stimuli were presented at 8 different orientations ranging from 0° to 315°. Stimuli were presented at the average preferred retinotopic location of the current field of view of axons which was determined using the above retinotopic mapping stimulus. Each grating stimulus was presented for 2 seconds with a 1 second interstimulus interval and each position was repeated 10 times in a pseudorandom order.

Stimuli to characterise spatial and temporal frequency selectivity consisted of 30 x 30° circular horizontally oriented square wave gratings presented at the average preferred retinotopic location of the current field of view of axons which was determined using the above retinotopic mapping stimulus, and with a fixed horizontal orientation. Additionally, a blank stimulus was presented. Spatial frequency was varied between 0.02 and 0.32 cycles per degree ({0.02, 0.04, 0.08, 0.16, 0.24, 0.32}) and temporal frequency between 0.5 and 24 Hz ({0.5, 2, 4, 8, 16, 24}) and each permutation of spatial and temporal frequency was presented for 2 seconds with a 1 second interstimulus interval and repeated 10 times in a pseudorandom order.

Stimuli to measure radial flow directional preference were constructed as video using a custom Python script. Stimuli occupied 50 degrees of visual space, and a spatial frequency of 0.1 cycles per degree and temporal frequency of 1 Hz. The stimulus appeared stationary for 5 seconds, flowed inwards (contraction) or outwards (dilation) for 5 seconds, and was then stationary for 5 seconds, followed by a 2 second inter-stimulus grey screen period. Each stimulus repeated 10 times in a pseudorandom order.

### Calcium imaging data processing

Calcium imaging data was registered and segregated into neuronal regions of interest (i.e. RSC boutons or V1 somas) using Suite2P (*25*). The time series of each ROI was then converted from a raw fluorescence value to ΔF/F with the denominator F value trace constructed by calculating the 5th percentile of the smoothed F value within a 20 second window centred on each sample in the F trace.

## Quantification and statistical analysis

### Analysis of retinotopic selectivity of RSC boutons and V1 neurons

In retinotopic mapping experiments, neurons/boutons were classified as visually responsive based on an ANOVA conducted across responses to all stimulus locations and the blank, with a significance threshold of p < 0.05 applied to determine responsiveness, as well as requiring that a bouton has a mean response to preferred stimulus which was greater than 2 standard deviations above the average blank response. Average response amplitude to stimulation in each position was then calculated for each visually responsive neuron/bouton with response calculated by averaging ΔF/F between 0.5 and 2 seconds after stimulus onset. To estimate retinotopic positional preference, a two-dimensional Gaussian function was fitted to the averaged responses at each retinotopic location using a least squares approach implemented in Python with the SciPy package. Fits with R^2^ < 0.7 were excluded from further analysis. The following function was fitted, in which A denotes the peak amplitude, (*x*0, *y*0) denotes the center of the Gaussian fit, and σ*x* and σ*y* correspond to the standard deviation of the function in the azimuth and elevation dimensions respectively:

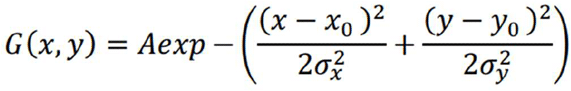

After fitting, the peak of the Gaussian distribution (*x*0, *y*0) was designated as the retinotopic preference, while the standard deviation in x and y (σ*x*, σ*y*) indicated the spatial extent of the receptive field in azimuth and elevation. In analysis in which positional preference differences were presented (either between each bouton and other boutons in the field of view or between each bouton and local V1 neurons) azimuth or elevation preference of the single bouton was subtracted from that of the larger reference population (i.e. other boutons or the V1 population). In azimuth, positive Δ position preference values thus indicate a bouton has a positional preference that is more nasal (i.e. towards the binocular visual field) than that of the reference population.

### Analysis of orientation tuning

In orientation tuning experiments, RSC boutons were classified as visually responsive based on an ANOVA conducted across responses to all stimulus orientations and the blank, with a significance threshold of p < 0.05 applied to determine responsiveness, as well as requiring that a bouton has a mean response to preferred stimulus which was greater than 2 standard deviations above the average blank response. Average response amplitude to stimulation in each orientation was then calculated for each visually responsive bouton with response calculated by averaging ΔF/F between 0.5 and 2 seconds after stimulus onset. Orientation selectivity was assessed in individual boutons by computing an Orientation Selectivity Index (OSI) using a vector-based method. This approach accounts for the circular nature of orientation tuning by summing neuronal responses as complex vectors in the polar coordinate system. The OSI was defined as:

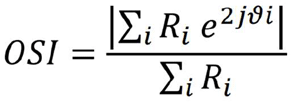

Where R_?_ denotes the bouton response to a stimulus presented at orientation θ_?_ (in radians), and j is the imaginary unit (j = *√*™1). The factor 2θ_?_ ensures periodicity over 180°, reflecting orientation rather than direction selectivity. The numerator represents the vector sum of orientation responses, which captures both the strength and angular bias of the preference. The denominator normalizes this by the total response magnitude across all orientations, constraining OSI values between 0 and 1. A higher OSI indicates stronger selectivity for a specific orientation, whereas an OSI near 0 reflects weak or absent selectivity.

### Analysis of spatial and temporal frequency tuning

In spatial and temporal frequency tuning experiments, RSC boutons or V1 neurons were classified as visually responsive based on an ANOVA conducted across responses to all stimulus spatial and temporal frequencies and the blank, with a significance threshold of p < 0.05 applied to determine responsiveness, as well as requiring that a bouton has a mean response to preferred stimulus which was greater than 2 standard deviations above the average blank response. Average response amplitude to each spatial and temporal frequency pair was then calculated for each visually responsive bouton/neuron with response calculated by averaging ΔF/F between 0.5 and 2 seconds after stimulus onset. To estimate spatial/temporal frequency preference, a two-dimensional Gaussian function was fitted to the averaged responses at for each spatial/temporal frequency pair using a least-squares approach implemented in Python with the SciPy package. The following two-dimensional Gaussian function was fitted, in which A denotes the amplitude at the preferred spatial/temporal frequency, *SF*_0_ and *TF*_0_ correspond to the preferred spatial and temporal frequencies of the bouton/neuron, and σ_SF_ and σ_TF_ correspond to the standard deviation of the function in the spatial and temporal frequency dimensions respectively:

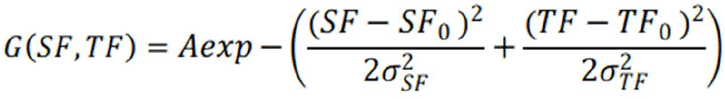

After fitting, the peak of the 2d Gaussian distribution (*SF*_0_ and *TF*_0_) was designated as the spatial/temporal frequency preference of each bouton/neuron, while the standard deviation in spatial/temporal frequency preference (σ_SF_ and σ_TF_) indicated selectivity.

### Other statistical methods

Statistical analyses were performed using Python (SciPy library), and group average values are presented throughout as mean ± standard error of the mean unless otherwise noted. The statistical significance of comparisons between groups was determined using a two-sided t test or ANOVA unless otherwise noted, and p values <0.05 were considered significant. Appropriateness of statistical tests used were determined by analysis of normality and homogeneity of variance using the Shapiro–Wilk and Brown–Forsythe tests. Precise group sizes were not decided in advance, but approximate group sizes were based on typical sizes used in this field in similar experiments.

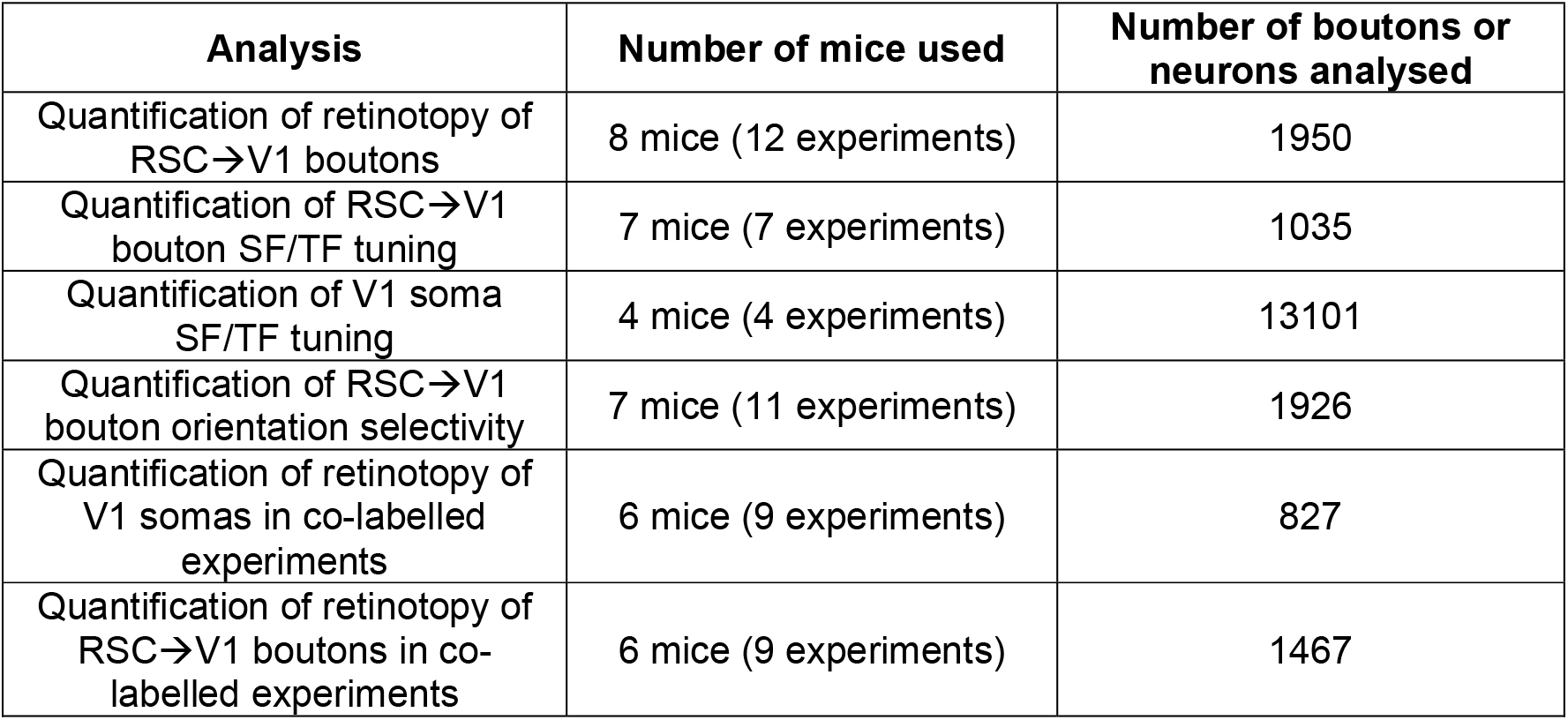

| **Analysis** | **Number of mice used** | **Number of boutons or neurons analysed** |
| --- | --- | --- |
| Quantification of retinotopy of RSC→V1 boutons | 8 mice (12 experiments) | 1950 |
| Quantification of RSC→V1 bouton SF/TF tuning | 7 mice (7 experiments) | 1035 |
| Quantification of V1 soma SF/TF tuning | 4 mice (4 experiments) | 13101 |
| Quantification of RSC→V1 bouton orientation selectivity | 7 mice (11 experiments) | 1926 |
| Quantification of retinotopy of V1 somas in co-labelled experiments | 6 mice (9 experiments) | 827 |
| Quantification of retinotopy of RSC→V1 boutons in co-labelled experiments | 6 mice (9 experiments) | 1467 |

## Notes

### Competing Interest Statement

The authors have declared no competing interest.

